# Reassortment Network of Influenza A Virus

**DOI:** 10.1101/2021.09.09.459621

**Authors:** Xingfei Gong, Mingda Hu, Wei Chen, Haoyi Yang, Boqian Wang, Junjie Yue, Yuan Jin, Long Liang, Hongguang Ren

## Abstract

Influenza A virus (IAV) genomes are composed of eight single-stranded RNA segments. Genetic exchange through reassortment of the segmented genomes often endows IAVs with new genetic characteristics, which may affect transmissibility and pathogenicity of the viruses. However, a comprehensive understanding of the reassortment history of IAVs remains lacking. To this end, we assembled 40296 whole-genome sequences of IAVs for analysis. Using a new clustering method based on Mean Pairwise Distances in the phylogenetic trees, we classified each segment of IAVs into clades. Correspondingly, reassortment events among IAVs were detected by checking the segment clade compositions of related genomes under specific environment factors and time period. We systematically identified 1927 possible reassortment events of IAVs and constructed their reassortment network. Interestingly, minimum spanning tree of the reassortment network reproved that swine act as an intermediate host in the reassortment history of IAVs between avian species and human. Moreover, reassortment patterns among related subtypes constructed in this study are consistent with previous studies. Taken together, our genome-wide reassorment analysis of all the IAVs offers an overview of the leaping evolution of the virus and a comprehensive network representing the relationships of IAVs.

**Availability:** Source code freely available for download at https://github.com/ElevenStr/IAVreassortment, implemented in python.

## 1 Introduction

Influenza A Virus (IAV) is a highly infectious viral pathogen that causes seasonal epidemics, occasional pandemics, and zoonotic outbreaks, which may lead to substantial human morbidity and mortality and a considerable financial burden worldwide (Liu, et al., 2020). It has been more than one hundred years since the Spanish flu (H1N1) virus caused the first recorded influenza pandemic, which is considered to be the most lethal natural event in modern history (Liu, et al., 2018; Smith, 2011). Since then, there have been three other pandemics caused by A(H2N2), A(H3N2), and A(H1N1)pdm09 viruses—the 1957 Asian flu, the 1968 Hong Kong flu, and the 2009 swine-origin flu, respectively. There are currently 18 HA (hemagglutinin) subtypes and 11 NA (neuraminidase) subtypes in IAVs, most of which spread in wild birds. Comparatively, only three combinations of HA and NA subtypes are known to be widespread in humans: H1N1, H2N2 and H3N2, of which H1N1 and H3N2 subtypes viruses cause seasonal epidemics (Petrova and Russell, 2018). The WHO estimated that seasonal influenza viruses infect 5-15% of the human population each year, causing approximately 500,000 deaths worldwide (Stohr, 2002).

As segmented RNA viruses, IAVs can exchange the gene segments through reassortment during co-infection. Specifically, when two or more IAVs infect the same cell, a hybrid virus can be produced by assembling the gene segments of the parental viruses into a nascent virion (McDonald, et al., 2016). Reassortment and mutation are both the main driving forces for the evolution of IAVs. However, the effect caused by mutations need to be accumulated for a long time, while the reassortment is often a leapfrog evolution. Reassortment also plays an important but unclear role in the emergence of the novel viruses and cross-species transmission (Dhanasekaran, et al., 2015). The virus that caused the 2009 H1N1 pandemic was generated from a triple-reassortment of H1N1 avian virus, H1N1 classical swine virus, and H3N2 human seasonal virus (Smith, et al., 2009). Furthermore, swine has been considered as the “mixing vessels” for IAVs since there are both α-2,3 and α-2,6 sialic acids (SAs) in their respiratory tracts, which provides conditions for the occurrence of reassortment (Ma, et al., 2009).

In recent years, a lot of studies for IAVs are based on mutations, especially in the receptor binding region on HA segment, while the study on reassortment detection method and the reassortment history of IAVs is relatively few. Rabadan et al. proposed a method to detect reassortment by comparing the sequence differences between two viruses (Rabadan, et al., 2008). This method considered that the number of differences in each segment sequence of two viruses should be proportional if there is no reassortment. Another algorithm was proposed by Silva et al. based on the neighborhood of strains (Silva, et al., 2012), which determined the reassortant virus by the size of the common neighborhood of two segments. Müller NF et al. used a coalescent-based model to study the reassortment pattern of different human influenza datasets and found that the reassortment rates of different human influenza viruses are very different (Muller, et al., 2020). These rare researches on IAVs reassortment tend to focus on viruses causing a single epidemic or with a single subtype, resulting in a lack of overall understanding of the reassortment history of IAVs. Ding X et al. collected the reassortment events of influenza A virus from published literature and constructed the FlueReassort database (Ding, et al., 2020). In our work, we proposed a novel genotype nomenclature for IAVs and used it to detect the reassortment in IAV genomes. The reassortment history and gene flow network were constructed, by which we found the characteristics and patterns in frequent reassortment events of IAVs.

## 2 Methods

### 2.1 Data preparation and processing

We assembled all the whole-genomes sequences of IAVs as of October 13, 2020 from the National Centre for Biotechnology Information (NCBI) website resources (https://ftp.ncbi.nih.gov/genomes/INFLUENZA/). After quality control, we obtained 40296 genomes of IAVs with complete sequence length and essential epidemiological information. Multiple sequence alignment for each segment was performed using MAFFT v7.037 (Katoh and Standley, 2013). To remove redundancy of sampling, we further filtered the genome sequences. Genome sequences with the same host, location, subtype, sampling year and sharing similarities over 99% were resampled. The relationships among represent strains and other strains were shown in Supplementary Table S1.

The phylogenetic tree was reconstructed for each segment by IQ-TREE (Nguyen, et al., 2015) using PhyloSuite platform (Zhang, et al., 2020) with the GTR+I+G4+F substitution model. Bat IAVs are used as outgroups to root the trees. We divided the hosts of these IAVs into human, swine and avian. The avian hosts were further divided into shorebirds, waterfowl, land birds and domestic birds. The avian host classifications were shown in Supplementary Table S2. The locations of the strains were mapped onto 22 areas according to the administrative division, including North America, Western Europe and East Asia et al (Supplementary Table S3).

### 2.2 Segment detailed type determination

In order to quantitatively divide the clade of each segment of the virus, we clustered the leaf nodes in the evolutionary tree based on the Mean Pairwise Distance (MPD). Therefore, sequences of each segment of IAVs were divided into different detailed types. MPD is the mean phylogenetic distance between the leaf nodes of an internal node on the phylogenetic tree. For example, the MPD of an internal node with n leaf nodes on PB1 segment phylogenetic tree is calculated as:

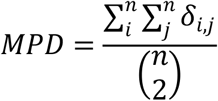

*δ*_*i,j*_ is the phylogenetic distance between leaf node i and leaf node j. 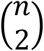 represents the combination number formula. The mean MPD of all internal nodes was used as a threshold to define the leaf nodes of an internal node as a specific cluster, namely a detailed type. We then assigned a index to each detailed type. For example, PB1_1 represents a detailed type of PB1 segment, consisting of a cluster of PB1 sequences.

### 2.3 Genotype nomenclature

In the work of Lu et al. (Lu, et al., 2007), the genotype was defined as the combination of the lineages for each segment in an IAV genome. In this paper, the genotype of an IAV genome was defined as the sequential combination of the detailed type for each segment. For example, [PB2_292, PB1_415, PA_333, HA_422, NP_463, NA_405, MP_369, NS_406] is the genotype of a H7N9 virus.

### 2.4 Reassortment definition and identification

A direct way to detect the reassortment of IAVs is to compare the positions on phylogenetic trees of different segments of the genome (Nelson, et al., 2008). Based on our genotype nomenclature, a reassortment was defined as a mixation or overlapping of the different genotype combinations. The mixation of different genotype combinations can be understood as the exchange of segments among different genotypes. If the genotype of a virus can be produced by the combination of the genotypes for two or more parental viruses and their epidemiological information is related, then we consider that this virus might be reassorted from these parental viruses. There may be more than one possible combination of parental viruses to generate the reassortant child. For each strain of each genotype, we screened all the genomes sampled within 5 years before the strain to find the most possible reassortment event with the least evolutionary cost, which was calculated as follows:

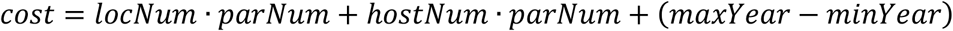

Where *locNum, hostNum, parNum* is the number of locations, hosts and parental viruses involved in the reassortment event. *maxYear* and *minYear* is the latest and the earliest year of the parental viruses. In this paper, only double-reassortment and triple-reassortment events are examined, as reassortment among more than three strain viruses is considered highly unlikely.

### 2.5 Gene flow network construction

A connecting network can be constructed using the genotypes as nodes and the relationships among genotypes as edges. We calculated two distance values, *dist* and *diff* between each pair of genotypes, where *dist* is the number of different segment detailed types between two genotypes, *diff* is the minimum difference of hosts, locations and sampling years combined among the viruses contained in the genotypes. A minimum spanning tree (MST), representing the gene flow network of IAVs, was established by traversing the network in pursuing the minimal reassortment cost. The MST was visualized using cystoscope v3.6.0 (Shannon, et al., 2003).

## 3 Results

### 3.1 Genotype definition for IAVs

We obtained 8932 high-quality and nonredundant representative IAV genomes from 40296 whole-genome database, where each segment of IAVs was divided into a detailed type. The segments of IAVs showed high diversity in Table 1, in which each segment was divided into hundreds of detailed types. As an example, we showed the detailed type determination result for PB1 segment in Figure 1. We listed the year range, hosts, locations, and subtypes where the detailed type emerged. The detailed type determination results for other segments were shown in Supplementary Figure SF1-7. After determining the detailed types for all segments of IAVs, we got the genotypes of all the IAV genomes, which consists of 6888 unique genotypes. In fact, these 6888 genotypes represent the genotypes of the previous 40296 viruses since each filtered virus can use the genotype of its representative virus as its genotype. The genotype of each virus was shown in Supplementary Table S4.

**Table 1.**
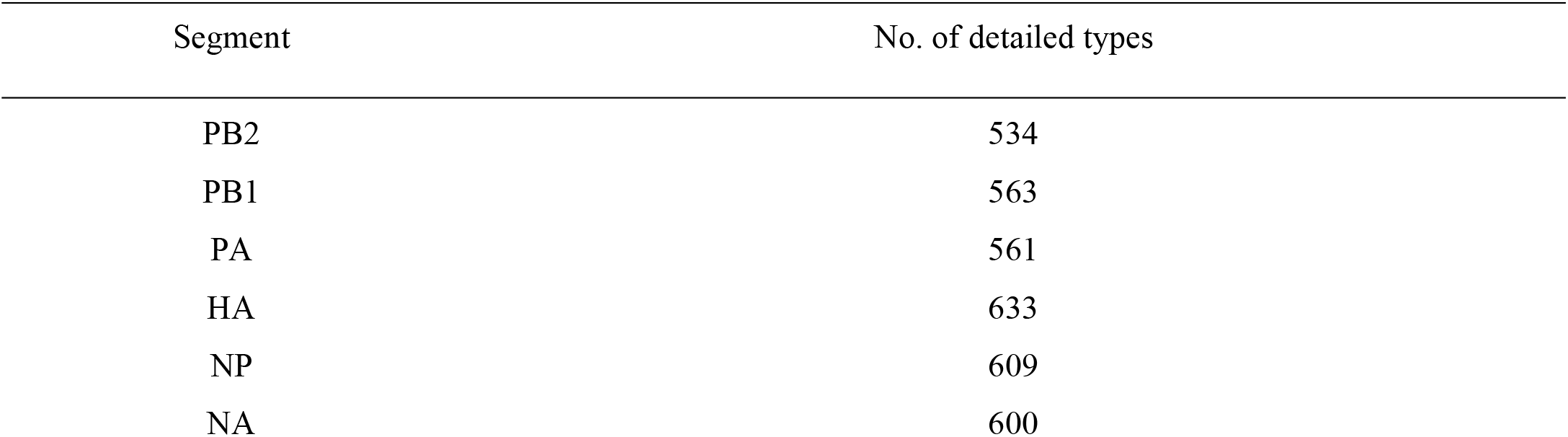
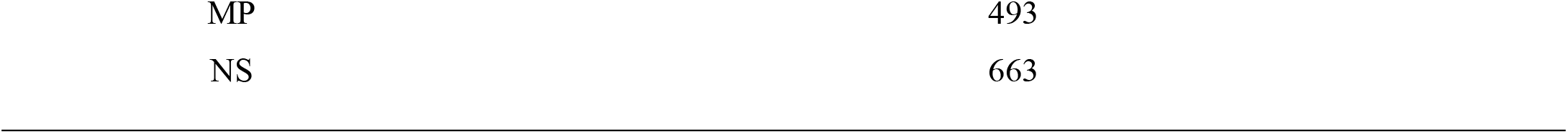
The number of detailed types for each segment.

| Segment | No. of detailed types |
| --- | --- |
| PB2 | 534 |
| PB1 | 563 |
| PA | 561 |
| HA | 633 |
| NP | 609 |
| NA | 600 |
| MP | 493 |
| NS | 663 |

**Figure. 1.**
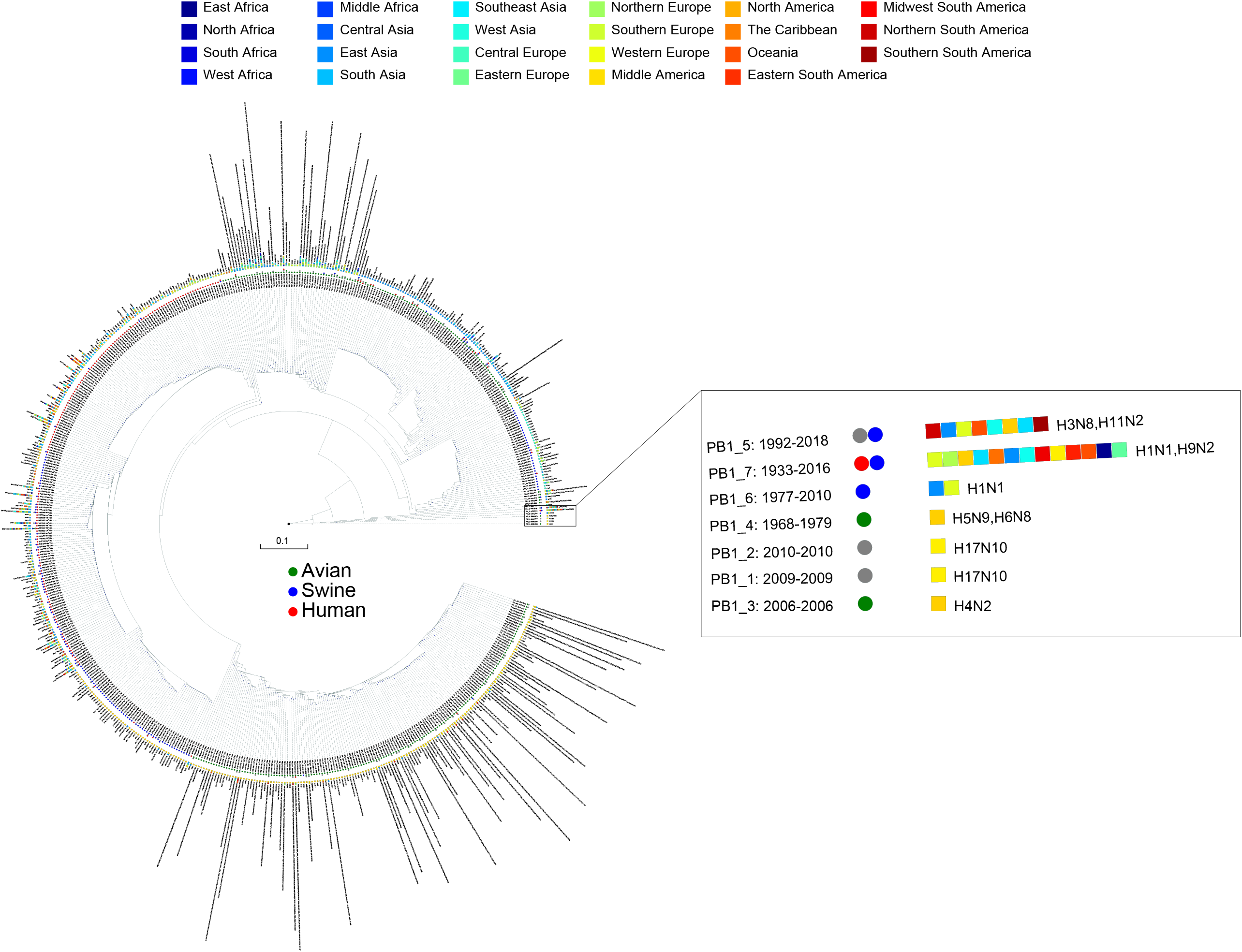
The detailed type result for PB1 segment. The year range, hosts, locations and subtypes are shown after each PB1 type, where the circles represent hosts and the rectangles represent locations. The hosts and locations are distinguished by different colors.

### 3.2 Reassortment detection

Combining the genotype and the epidemiological information of the virus, we immediately knew a segment detailed type had occurred in which year, which host, which location and which subtype. Meanwhile, for each segment of the IAV to be analyzed, we also knew which virus had the same segment detailed type. These viruses with identical segment detailed types may provide the reassortment source for the virus in analysis. In particular, we showed the result of reassortment analysis for the virus (A/WuXi/0409/2014) in Figure 2, where the viruses with identical segment detailed types were listed. The genotype of this H7N9 virus is [PB2_292, PB1_415, PA_333, HA_422, NP_463, NA_405, MP_369, NS_406]. Each segment detailed type had spread widely before the virus emerged, indicating that the virus had a high possibility to be a reassortant virus. Later we attempted to identify the reassortment source and found that the virus was might generated from a triple-reassortment of H7N9 virus (A/chicken/Huzhou/3791/2013), H9N2 virus (A/chicken/Suzhou/4954/2013) and H9N2 virus (A/chicken/Jiangsu/SIC11/2013) (Figure 3). The triple reassortant virus acquired its HA and NA segments from H7N9 virus (A/chicken/Huzhou/3791/2013). The PB2, PA, NP and MP segments were derived from H9N2 virus (A/chicken/Suzhou/4954/2013). Another H9N2 virus (A/chicken/Jiangsu/SIC11/2013) provided PB1 and NS segments for the triple reassortant virus. In a word, the H7N9 virus (A/WuXi/0409/2014) obtained HA and NA segments from its parental H7N9 virus, and internal segments from its parental H9N2 viruses. Obviously, with our segment detailed type determination and genotype nomenclature, it will be an easy task to detect reassortment for IAVs.

**Figure. 2.**
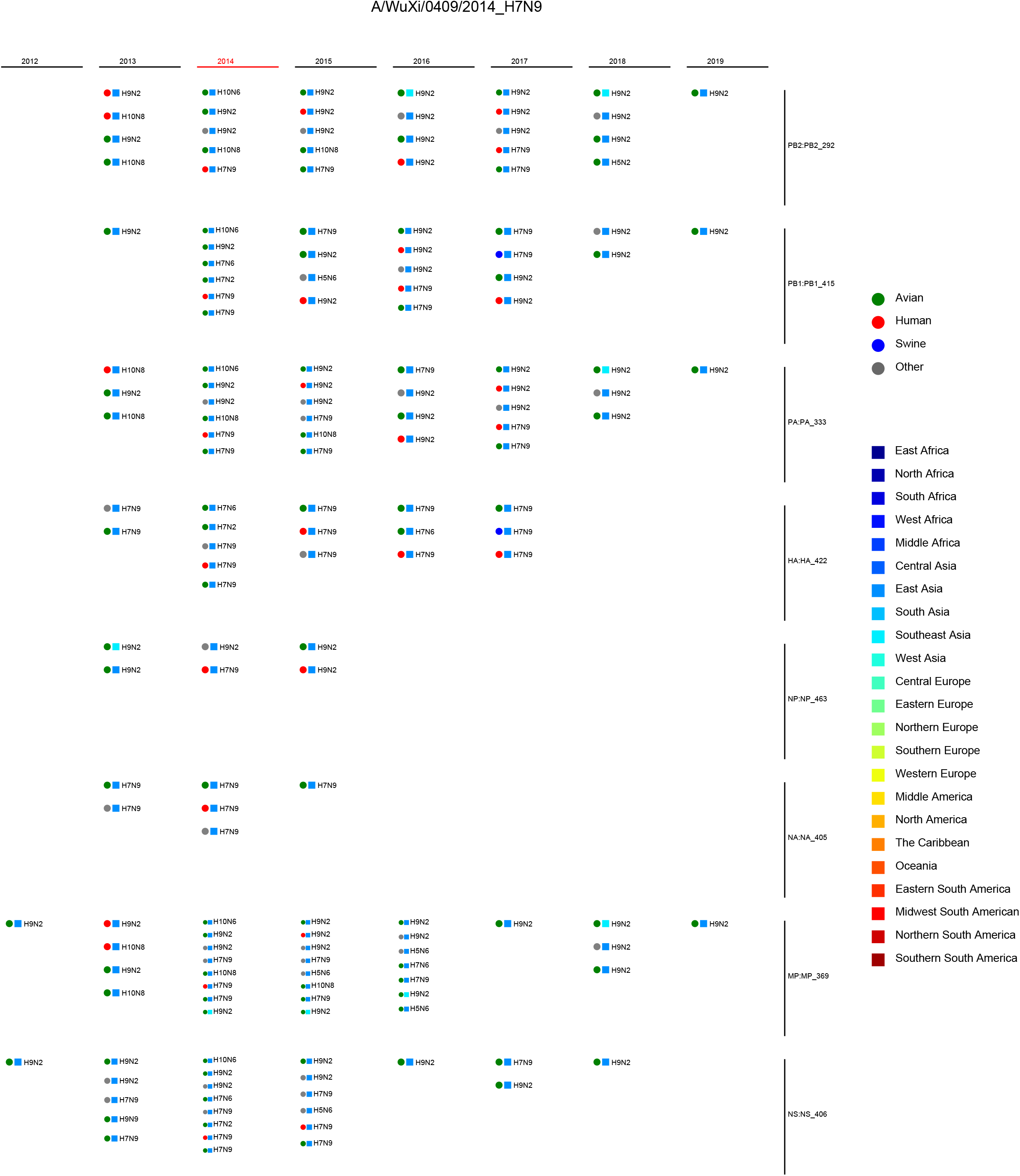
Reassortment analysis for the virus (A/WuXi/0409/2014). The genotype of the virus is [PB2_292, PB1_415, PA_333, HA_422, NP_463, NA_405, MP_369, NS_406]. The infor-mation of the viruses with same segment detailed types is shown, including host, location, subtype and year. The circles represent hosts and the rectangles represent locations, both of which are distinguished by different colors.

**Figure. 3.**
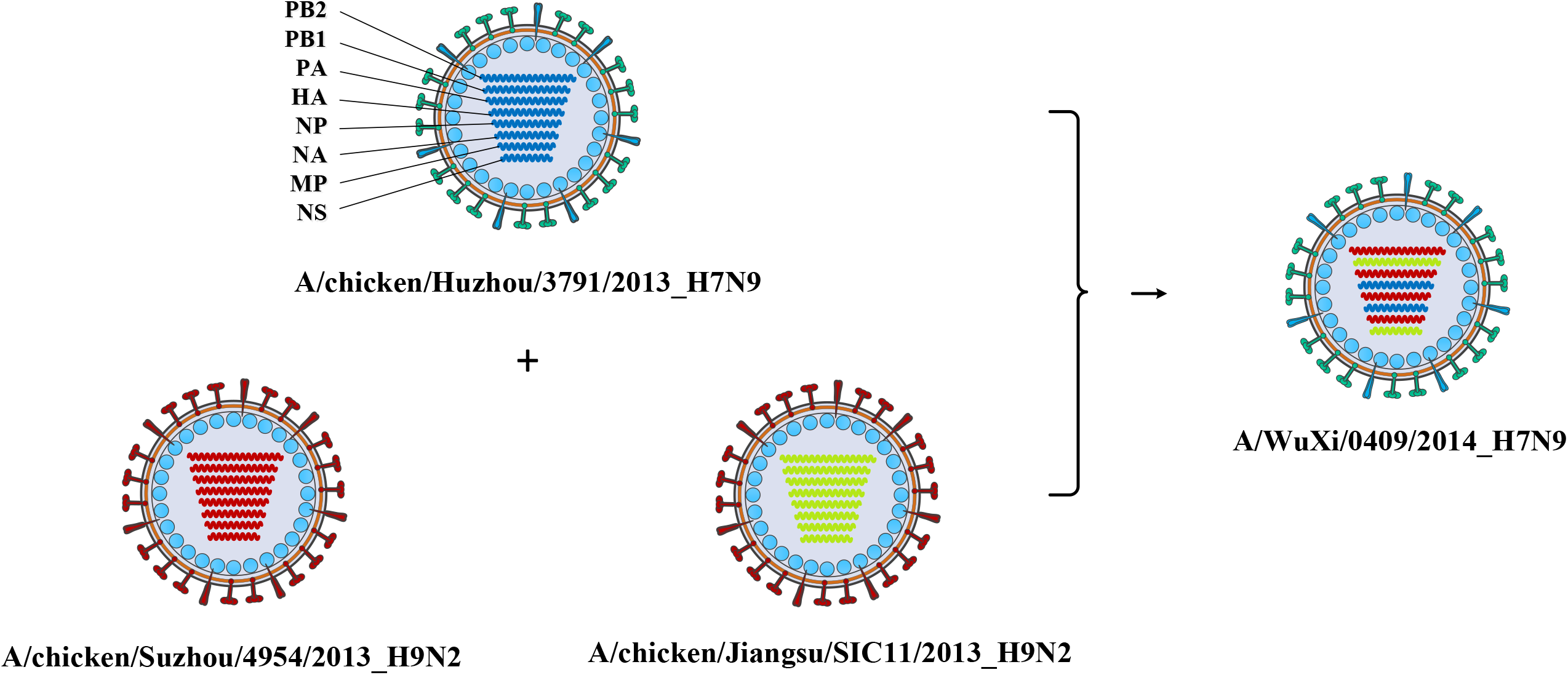
The reassortment source of the virus (A/WuXi/0409/2014). The virus was produced by a triple-reassortment of the virus (A/chicken/Suzhou/4954/2013), the virus (A/chicken/Huzhou/3791/2013) and the virus (A/chicken/Jiangsu/SIC11/2013). The genotype of the virus (A/chicken/Huzhou/3791/2013) is [PB2_309, PB1_396, PA_284, HA_422, NP_561, NA_405, MP_467, NS_406], which is indicated by blue. The genotype of the virus (A/chicken/Suzhou/4954/2013) is [PB2_292, PB1_252, PA_333, HA_523, NP_463, NA_285, MP_369, NS_387], which is indicated by red. The genotype of the virus (A/chicken/Jiangsu/SIC11/2013) is [PB2_310, PB1_415, PA_307, HA_536, NP_545, NA_385, MP_467, NS_406], which is indicated by green. The reassortant virus obtained HA and NA segments from a H7N9 virus, and internal segments from two H9N2 viruses.

### 3.3 The reassortment history of IAVs

In order to construct the reassortment history of IAVs, we detected the reassortment events for all viruses in our dataset, using the method we proposed above. The result of reassortment history was shown in Figure 4, in which we displayed each reassortment event as the connections from the parental viruses to the reassortant strain. We also shown another form of the result in Supplementary Figure SF8. It should be noted that the reassortment event we detected first emerged in North America, but this does not mean that the origin of IAVs was in North American, since the early sequencing technology around the world was not as advanced as it is now. The reassortment history of IAVs generally reflects the reassortant trend, in which we can know which locations and which hosts are more likely to generate reassortment. In our work, we found that the frequency of inter-locations reassortment is high in various locations of Asia and Europe, which can be illustrated from the large number of red lines in these locations in Figure 4. Although the number of reassortment events in North America is large, most of them are intra-locations reassortment. This finding was also reflected in Figure 5, in which we counted the number of reassortment events in different locations each year.

**Figure. 4.**
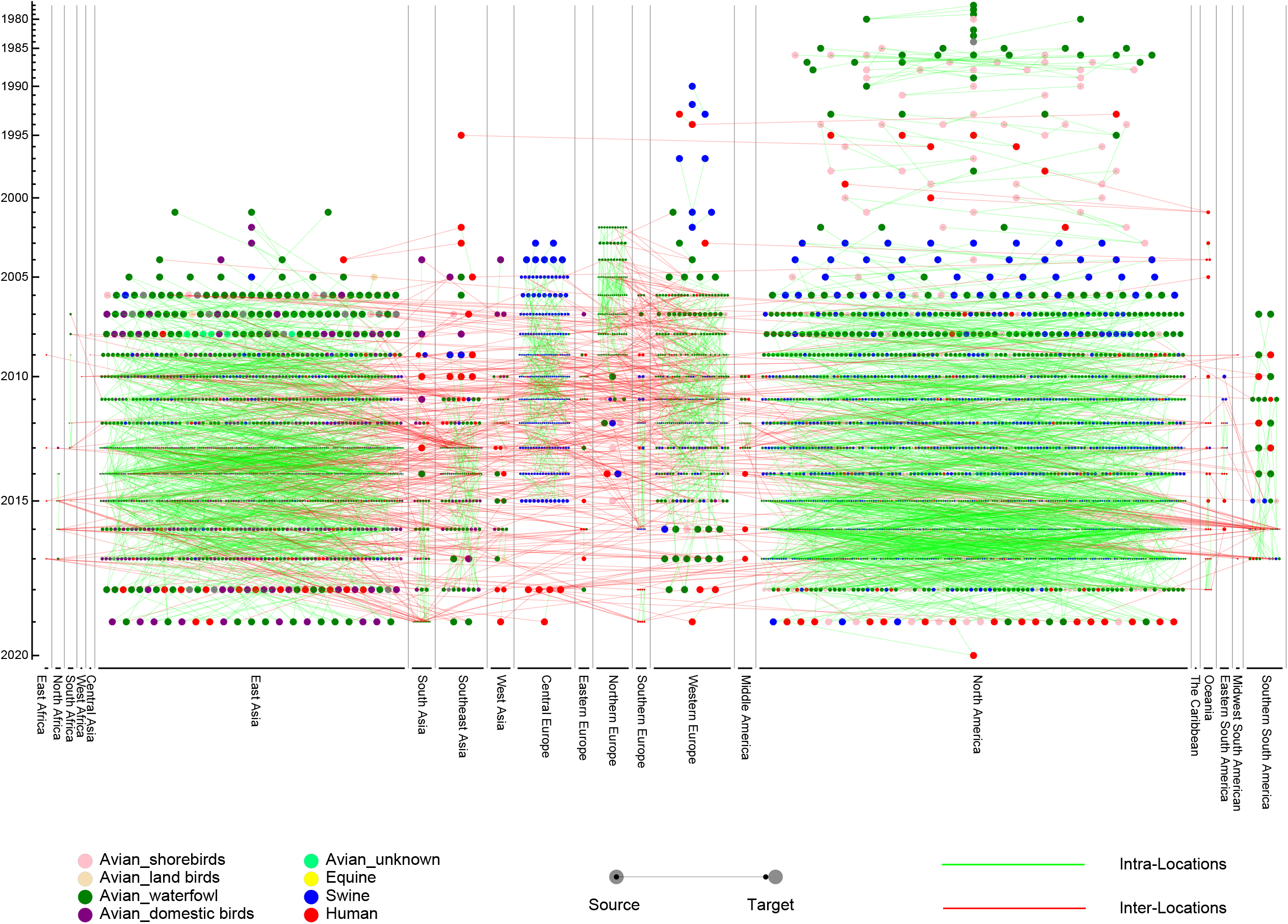
The reassortment history of IAVs. The horizontal axis represents the locations, while the vertical axis represents the years. Each circle represents a virus, with different colors to indicate the hosts. The line from source to target represents the parental virus produces the reassortant virus. Intra-Locations and Inter-Locations reassortment are indicated by green and red lines, respectively.

**Figure. 5.**
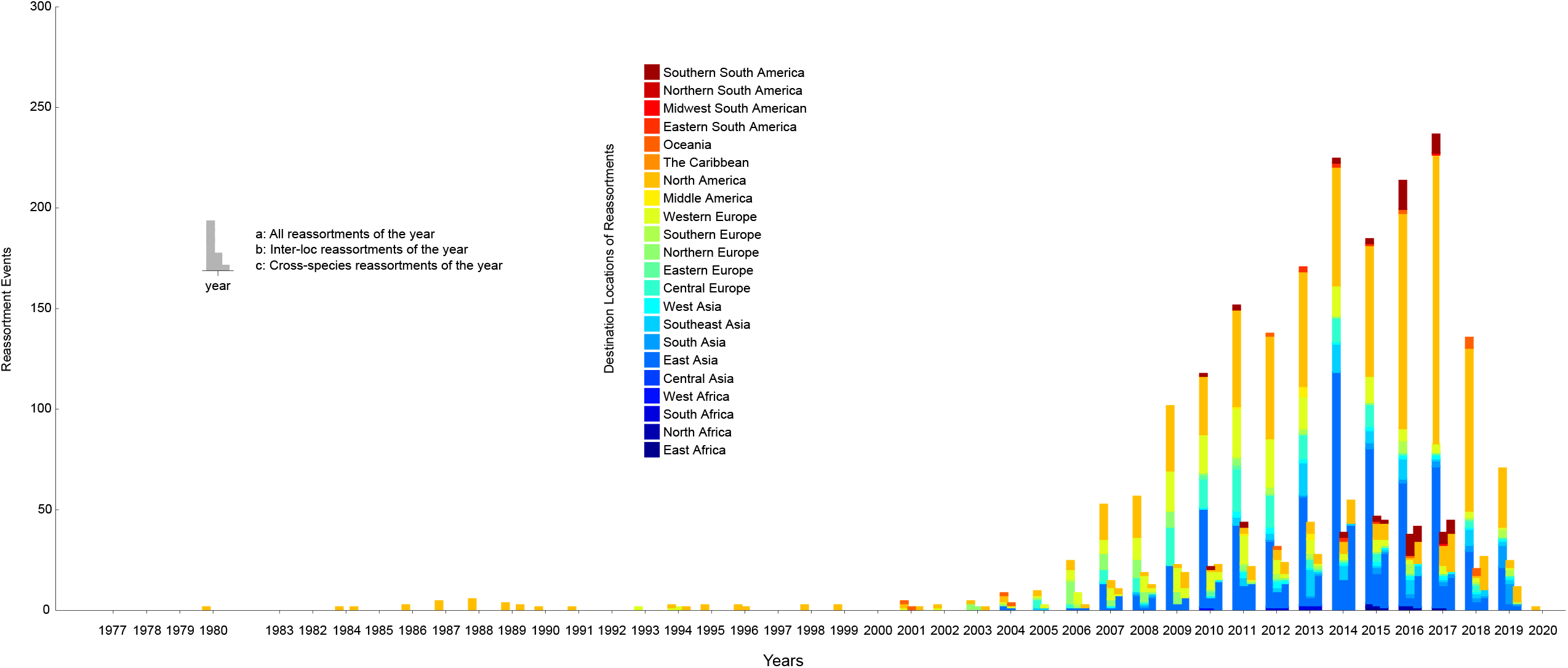
The number of reassortment events per year in different locations. The horizontal axis represents the years, and the vertical axis represents the number of reassortment events. The locations of the reassortant viruses are distinguished by different colors. The number of reassortment events per year was divided into three types for statistics: all reassortments, inter-location reassortments and cross-species reassortments.

### 3.4 Swine act as an intermediate host in the reassorment history of IAVs

In reassortment history of IAVs above, we limited the number of reassortment sources due to the limitation of calculation. Actually, the reassortment of IAVs is complicated, since the genome of an IAV can come from eight different viruses at most. In order to analyze how the genes of IAVs are transmitted during the reassortment, we constructed the gene flow network as shown in Figure 6. The network was composed of nodes and edges, where each node was a genotype of IAVs and the edge represented gene segment flow by reassortment between two genotypes. It should be noted that we found some conclusions which is consistent with previous studies after we added the host information to the network. Swine as an intermediate host played an important role in the gene flow between avian and human IAVs, which can be seen from the swine host nodes in the figure as the hub connecting the avian host nodes and the human host nodes. Actually, it is universally acknowledged that swine are the “mixed vessels” for IAVs since they can be infected by both avian and human IAVs and facilitate the reassortment events of the IAVs (Amélie, et al., 2018). Meanwhile, due to the global live swine trade, swine also plays a crucial role in the global spread of IAVs, which provides the possibility of cross-locations reassortment for IAVs (Nelson, et al., 2015). Therefore, our result on swine as an intermediate host is consistent with previous studies on the role of swine.

**Figure. 6.**
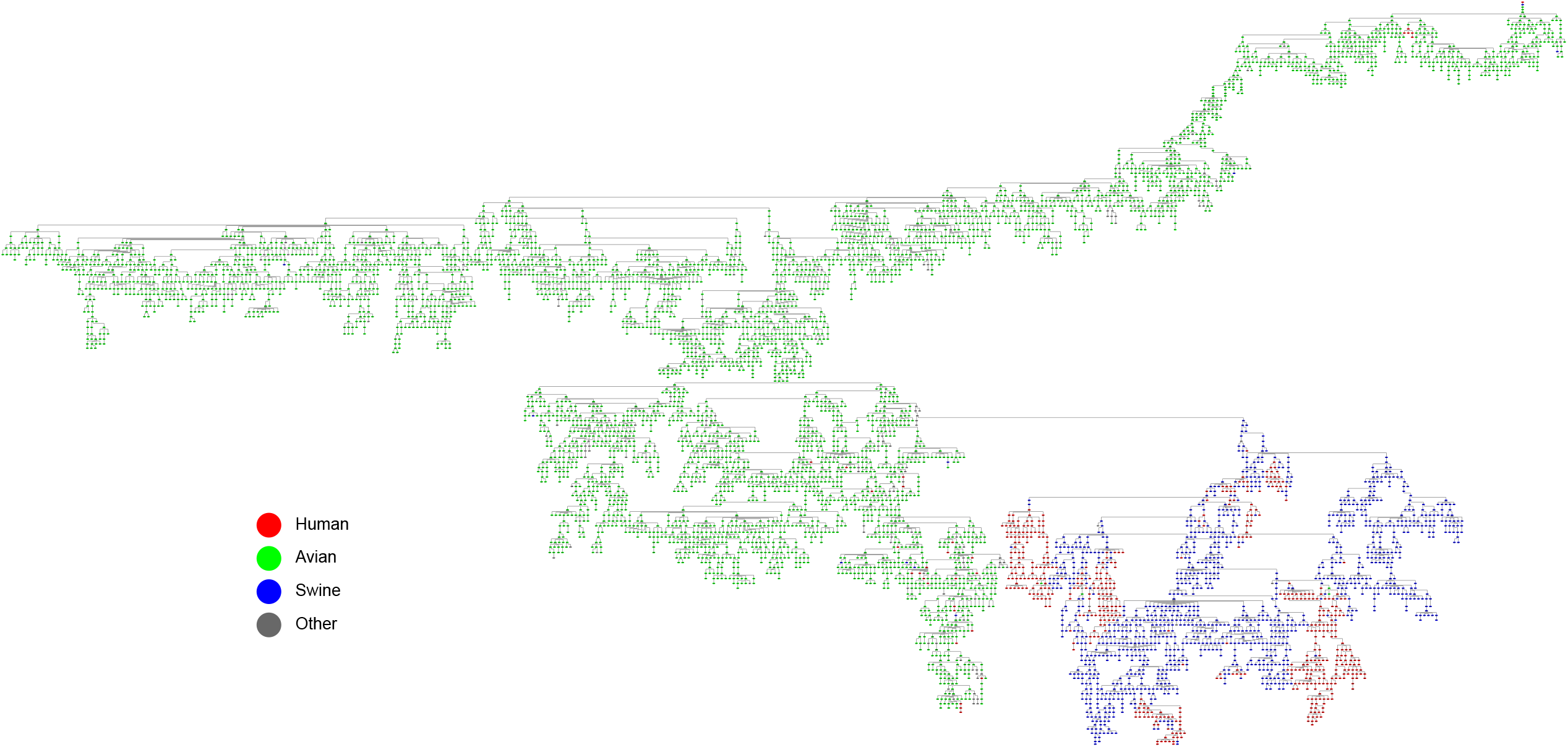
The gene flow network of IAVs (with host information annotated). Each node is a genotype of IAVs and the edge represents gene segment flow by reassortment between two genotypes. The hosts are distinguished by different colors, where red represents human, green represents avian, blue represents swine, and grey represent other hosts.

### 3.5 Frequent reassortment between H9N2 and H7N9 viruses

Analyzing the subtypes of IAVs reassortment can also help us understand the patterns of the reassortment. The avian influenza viruses with H9N2 subtype are widely distributed in different regions of China, and provide internal segments for IAVs for other subtypes, which poses a serious threat to public health (Liu, et al., 2014). In our study, we also found a similar result after we added the subtype information to the gene flow network. It has been shown in Supplementary Figure SF9 that there are many intersections between H9N2 and H7N9 viruses, which means that frequent reassortments occurred in IAVs with such subtypes. Later we found that almost all of these viruses originated from domestic birds in East Asia (Supplementary Figure SF10). It has been demonstrated that H9N2 has replaced H5N6 and H7N9 as the dominant AIV subtype in both chickens and ducks of China, and the novel viruses generated from reassortment continued to appear and spread among the avian hosts (Bi, et al., 2020). The internal segments of some H9N2 viruses have acquired fitness in human, while viruses with other subtypes such as H7N9 can acquire this fitness through reassortment with them (Liu, et al., 2013; Lu, et al., 2014; Zhang, et al., 2013). The reassortment analysis for the H7N9 virus (A/WuXi/0409/2014) (Figure 3) revealed that the virus obtained HA and NA segments from its parental H7N9 virus and internal segments from its parental H9N2 viruses. It is noteworthy that this reassortant virus was sampled from a human and its parental viruses originated from domestic birds. In other words, the parental viruses completed cross-species transmission through this reassortment patter.

## 4 Discussion

In order to verify the correctness of our clustering method, we used the unified nomenclature system for the highly pathogenic H5N1 avian influenza viruses (Organization, 2011) to cluster HA segments of H5N1 viruses in the database and compared with our result. The unified nomenclature system used three specific clade definition criteria: sharing of a common (clade-defining) node in the phylogenetic tree; monophyletic grouping with a bootstrap value of ≥60 at the clade-defining node; and average percentage pairwise nucleotide distances between and within clades of >1.5% and <1.5%. We marked the clustering results of the two methods on the phylogenetic tree, which was shown in Supplementary Figure SF11. The inner circle was the clustering result of the unified nomenclature system, and the outer circle was our clustering result. The clustering result of the unified nomenclature system was more detailed than ours, but the clustering boundaries of the two methods were mostly synchronized.

IAVs are rapidly evolving through reassortment, posing a huge challenge to vaccine development and a serious threat to public health. Traditional studies in IAVs reassortment often focus on the specific virus that causes a certain epidemic, lacking a systematic understanding of IAVs reassortment history. In the present study, we divided each segment of IAVs into detailed types according to the Mean Pairwise Distance (MPD) on the phylogenetic trees. A new method was proposed to define the genotype of IAVs, and the genotype nomenclature was then exploited to analyze whole genomes of IAVs and detect the reassortment. Generally, all the segments of the virus should be compared comprehensively when detecting the reassortment source of IAVs, rather than considering only a single segment. If some segments of a virus have high homology with a parental virus and low homology with another parental virus, and the circumstance in other segments is opposite, the virus is likely to be generated from a reassortment of the two parental viruses (Tian, et al., 2011). The reassortment detection method in our work not only compared the source of each segment but also considered the epidemiological information of the IAVs, which is more reasonable. Different from previous reassortment detection method (Niranjan and Carl, 2011; Suzuki, 2010; Svinti, et al., 2013; Yurovsky and Moret, 2011), we quantitatively represent the inconsistency of the phylogenetic trees as different segment detailed types. Meanwhile, the results produced by our method are consistent with previous studies, which confirms the effectiveness of our method. Therefore, the method we proposed can indeed effectively define the genotype of IAV and detect the reassortment, especially for large-scale data sets.

It should be pointed out that our result only listed the reassortment events we found. Some IAVs were generated from reassortment but we did not detect them because their parental viruses could not be found due to insufficient sampling or incomplete information. Our research was based on the sequence data and the epidemiological information, which benefits from the continuous monitoring and sequencing of IAVs by researchers around the world.

Taken together, we proposed a novel reassortment detection method based on our genotype nomenclature and systematically studied the history and patterns of reassortment in IAVs, which will be beneficial for the prevention and control of IAVs worldwide.

## References

Amélie, et al. Spatio-temporal distribution and evolution of the A/H1N1 2009 pandemic virus in pigs in France from 2009 to 2017: identification of a potential swine-specific lineage. Journal of virology 2018.

Bi, Y., et al. Dominant subtype switch in avian influenza viruses during 2016–2019 in China. Nature Communications 2020;11(1):5909.

Dhanasekaran, V., Mukerji, R. and Smith, G. RNA Virus Reassortment: An Evolutionary Mechanism for Host Jumps and Immune Evasion. PLoS pathogens 2015;11:e1004902.

Ding, X., et al. FluReassort: a database for the study of genomic reassortments among influenza viruses. Brief Bioinform 2020;21(6):2126–2132.

Katoh, K. and Standley, D. MAFFT Multiple Sequence Alignment Software Version 7: Improvements in performance and usability. Molecular biology and evolution 2013;30.

Liu, D., Shi, W. and Gao, G.F. Poultry carrying H9N2 act as incubators for novel human avian influenza viruses. Lancet 2014;383(9920):869–869.

Liu, D., et al. Origin and diversity of novel avian influenza A H7N9 viruses causing human infection: phylogenetic, structural, and coalescent analyses. The Lancet 2013;381(9881):1926–1932.

Liu, W., et al. On the Centenary of the Spanish Flu: Being Prepared for the Next Pandemic. Virologica Sinica 2018;33.

Liu, W.J., et al. Emerging HxNy Influenza A Viruses. Cold Spring Harbor Perspectives in Medicine 2020.

Lu, G., et al. FluGenome: a web tool for genotyping influenza A virus. Nucleic Acids Research 2007;35(Web Server):W275-W279.

Lu, J., et al. Continuing Reassortment Leads to the Genetic Diversity of Influenza Virus H7N9 in Guangdong, China. Journal of Virology 2014;88(15):8297–8306.

Ma, W., Kahn, R.E. and Richt, J.A. The pig as a mixing vessel for influenza viruses: Human and veterinary implications. Journal of Molecular & Genetic Medicine An International Journal of Biomedical Research 2009;3(1):158–166.

McDonald, S.M., et al. Reassortment in segmented RNA viruses: mechanisms and outcomes. Nature Reviews Microbiology 2016;14(7):448–460.

Muller, N.F., et al. Bayesian inference of reassortment networks reveals fitness benefits of reassortment in human influenza viruses. Proc Natl Acad Sci U S A 2020;117(29):17104–17111.

Nelson, M.I., et al. Multiple Reassortment Events in the Evolutionary History of H1N1 Influenza A Virus Since 1918. Plos Pathogens 2008;4(2):e1000012.

Nelson, M.I., et al. Global migration of influenza A viruses in swine. Nature Communications 2015;6(1):6696.

Nguyen, L.-T., et al. IQ-TREE: A Fast and Effective Stochastic Algorithm for Estimating Maximum-Likelihood Phylogenies. Molecular Biology and Evolution 2015;32(1):268–274.

Niranjan, N. and Carl, K. GiRaF: robust, computational identification of influenza reassortments via graph mining. Nucleic Acids Research 2011(6):e34.

Organization, W.H. Updated unified nomenclature system for the highly pathogenic H5N1 avian influenza viruses. Influenza 2011.

Petrova, V.N. and Russell, C.A. The evolution of seasonal influenza viruses. Nature Reviews Microbiology 2018;16(1):47–60.

Rabadan, R., Levine, A.J. and Krasnitz, M. Non - random reassortment in human influenza A viruses. Influenza and Other Respiratory Viruses 2008;2.

Shannon, et al. Cytoscape: A Software Environment for Integrated Models of Biomolecular Interaction Networks. Genome Research 2003;13(11):2498–2504.

Silva, U., et al. A comprehensive analysis of reassortment in influenza A virus. Biology open 2012;1:385–390.

Smith, G.J.D. The emergence of pandemic influenza viruses. International Journal of Infectious Diseases 2011;15:S7–S8.

Smith, G.J.D., et al. Origins and evolutionary genomics of the 2009 swine-origin H1N1 influenza A epidemic. Nature 2009;459(7250):1122–1125.

Stohr, K. Influenza-WHO cares. The Lancet Infectious Diseases 2002;2.

Suzuki, Y. A phylogenetic approach to detecting reassortments in viruses with segmented genomes. Gene 2010;464(1):11–16.

Svinti, V., Cotton, J.A. and McInerney, J.O. New approaches for unravelling reassortment pathways. BMC Evolutionary Biology 2013;13(1):1.

Tian, D., Wang, Y. and Tao, Z. A novel strategy for exploring the reassortment origins of newly emerging influenza virus. Bioinformation 2011;7(2):64–68.

Yurovsky, A. and Moret, B. FluReF, an automated flu virus reassortment finder based on phylogenetic trees. Bmc Genomics 2011;12(Suppl 2):S3–S3.

Zhang, D., et al. PhyloSuite: An integrated and scalable desktop platform for streamlined molecular sequence data management and evolutionary phylogenetics studies. Molecular Ecology Resources 2020;20(1):348–355.

Zhang, L., Zhang, Z. and Weng, Z. Rapid Reassortment of Internal Genes in Avian Influenza A(H7N9) Virus. Clinical infectious diseases : an official publication of the Infectious Diseases Society of America 2013;57.

