## Supplementary Figures for "Reassortment Network of Influenza A Virus": Supplementary Figure SF1.pdf

Supplementary Figure SF1. The detailed type result for PB2 segment. The year range, hosts, locations and subtypes are shown after each PB2 type, where the circles represent hosts and the rectangles represent locations. The hosts and locations are distinguished by different colors.

| | | | | | |
|---|---|---|---|---|---|
| East Africa | Middle Africa | Southeast Asia | Northern Europe | North America | Midwest South American |
| North Africa | Central Asia | West Asia | Southern Europe | The Caribbean | Northern South America |
| South Africa | East Asia | Central Europe | Western Europe | Oceania | Southern South America |
| West Africa | South Asia | Eastern Europe | Middle America | Eastern South America | |

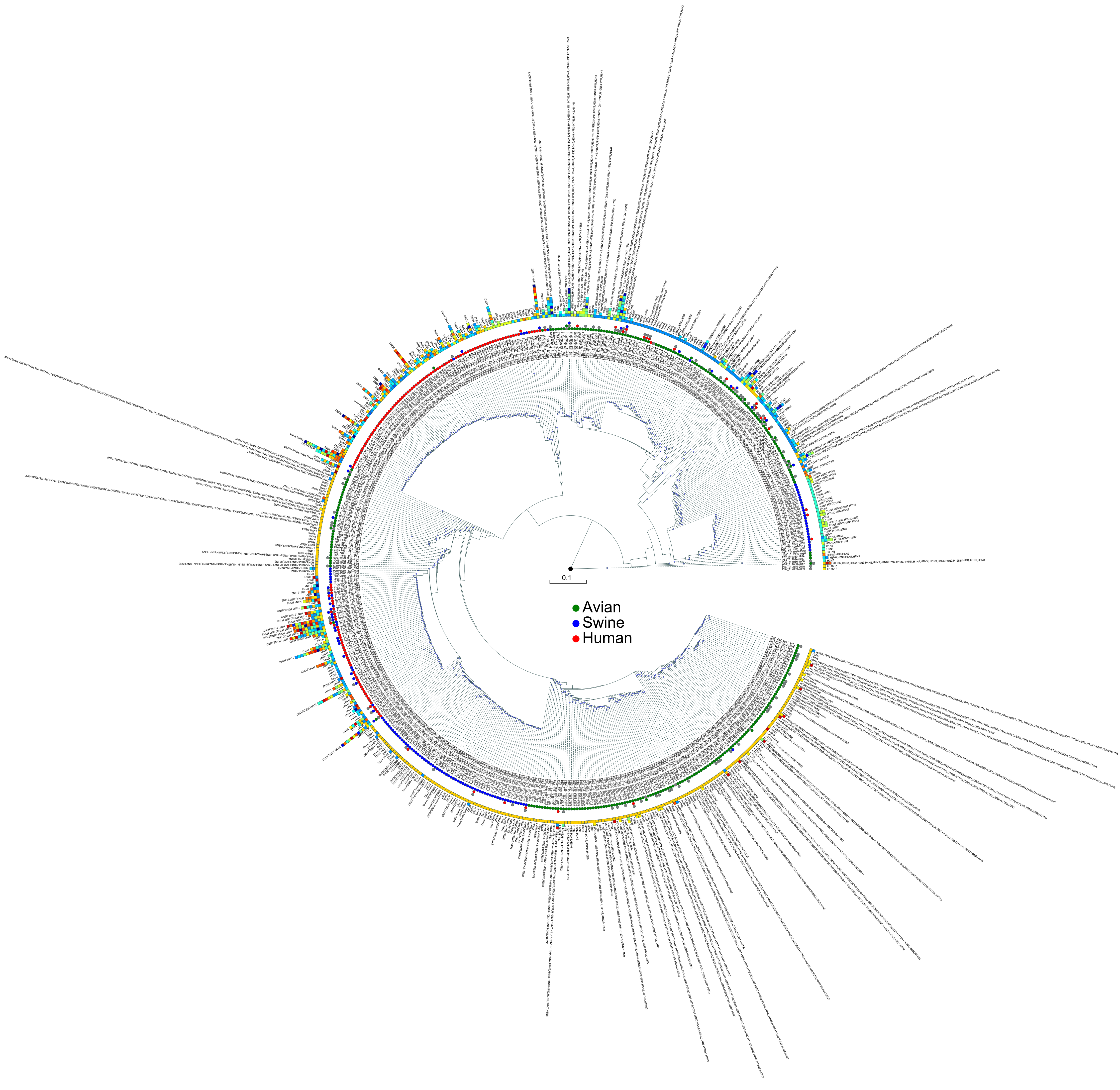
