## Supplementary Figures for "Reassortment Network of Influenza A Virus": Supplementary Figure SF2.pdf

Supplementary Figure SF2. The detailed type result for PA segment. The year range, hosts, locations and subtypes are shown after each PA type, where the circles represent hosts and the rectangles represent locations. The hosts and locations are distinguished by different colors.

East Africa
North Africa
South Africa
West Africa
Middle Africa
Central Asia
East Asia
South Asia
Southeast Asia
West Asia
Central Europe
Eastern Europe
Northern Europe
Southern Europe
Western Europe
Middle America
North America
The Caribbean
Oceania
Eastern South America
Midwest South American
Northern South America
Southern South America

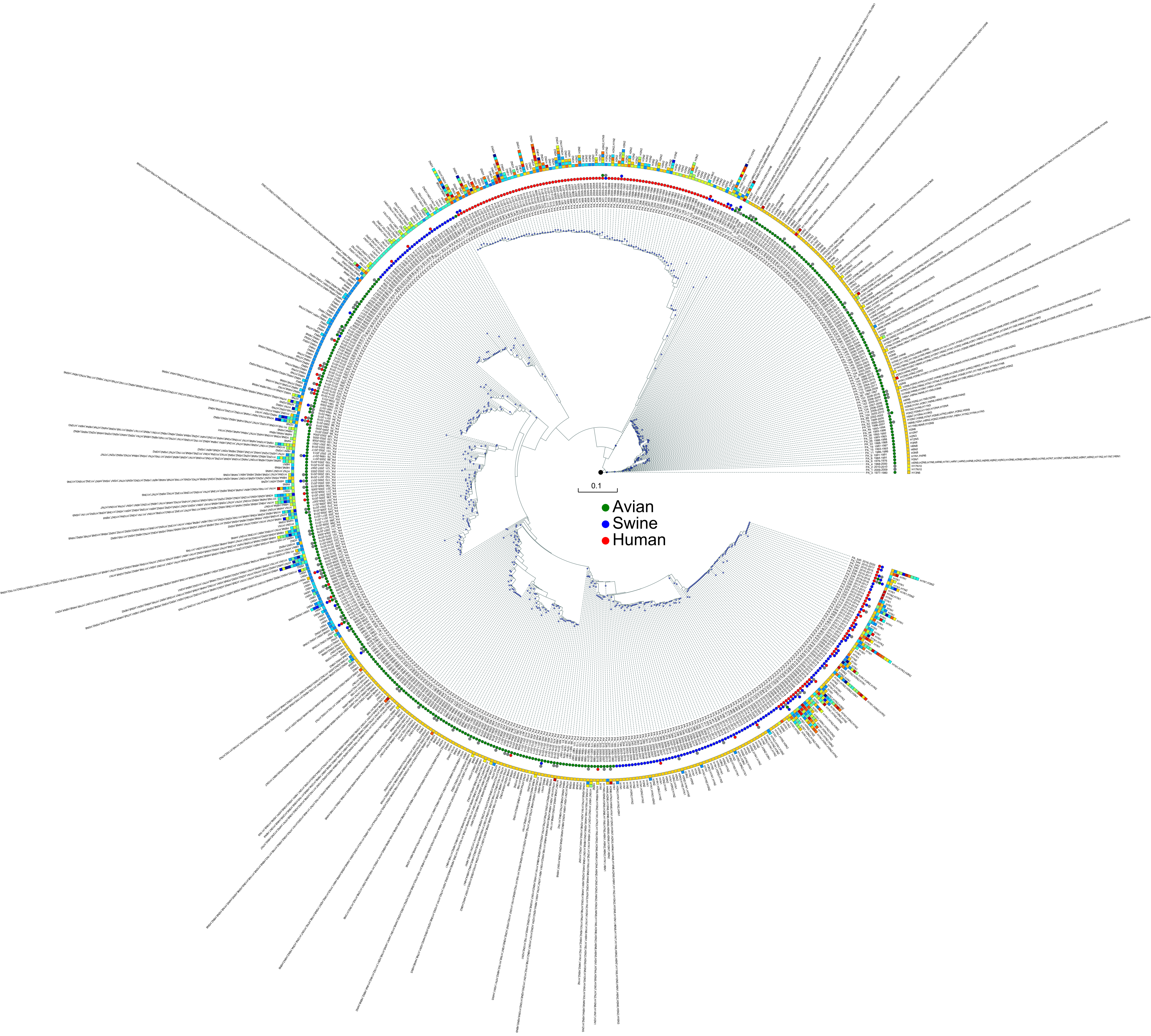
