## Supplementary Figures for "Reassortment Network of Influenza A Virus": Supplementary Figure SF3.pdf

Supplementary Figure SF3. The detailed type result for HA segment. The year range, hosts, locations and subtypes are shown after each HA type, where the circles represent hosts and the rectangles represent locations. The hosts and locations are distinguished by different colors.

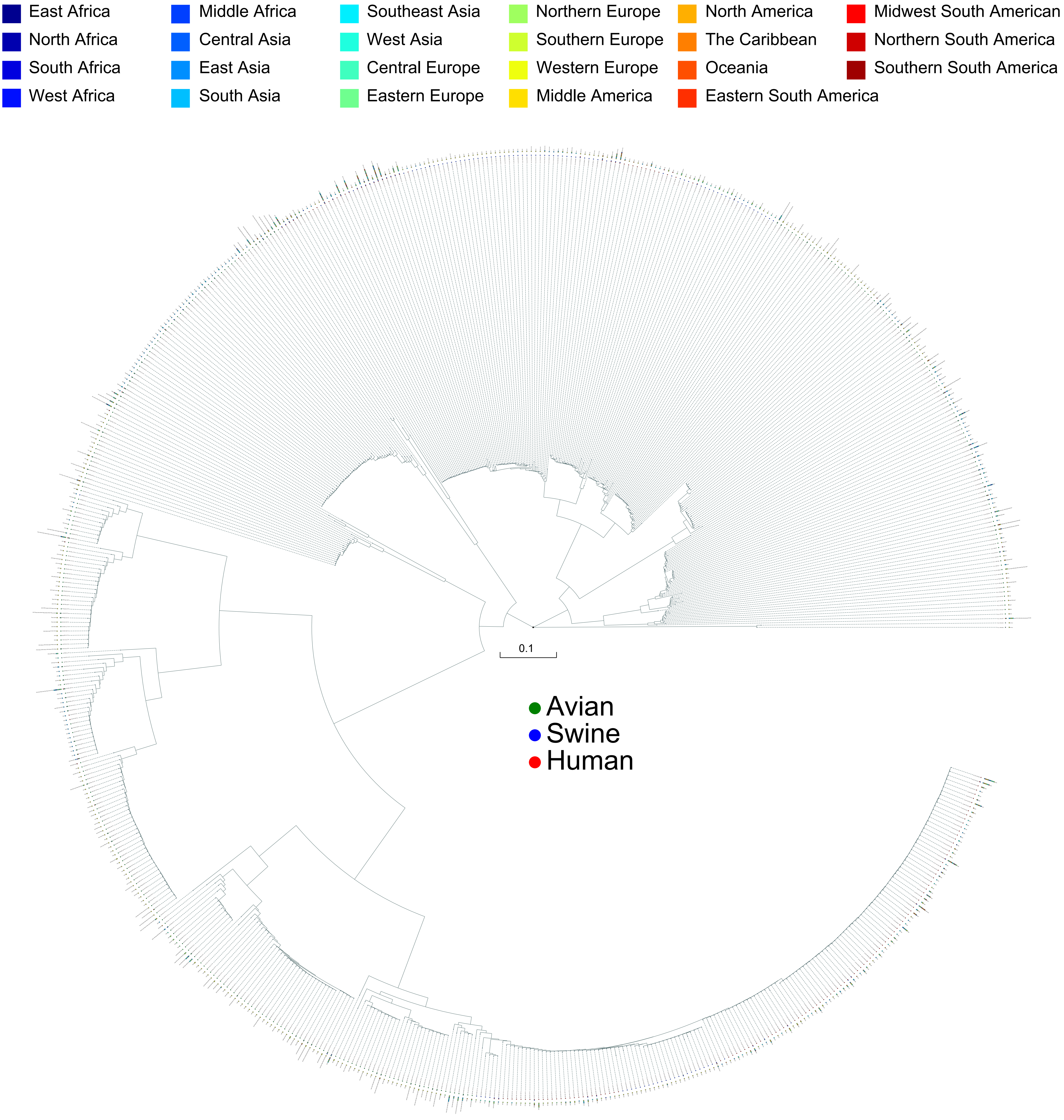
