## Supplementary Figures for "Reassortment Network of Influenza A Virus": Supplementary Figure SF4.pdf

Supplementary Figure SF4. The detailed type result for NP segment. The year range, hosts, locations and subtypes are shown after each NP type, where the circles represent hosts and the rectangles represent locations. The hosts and locations are distinguished by different colors.

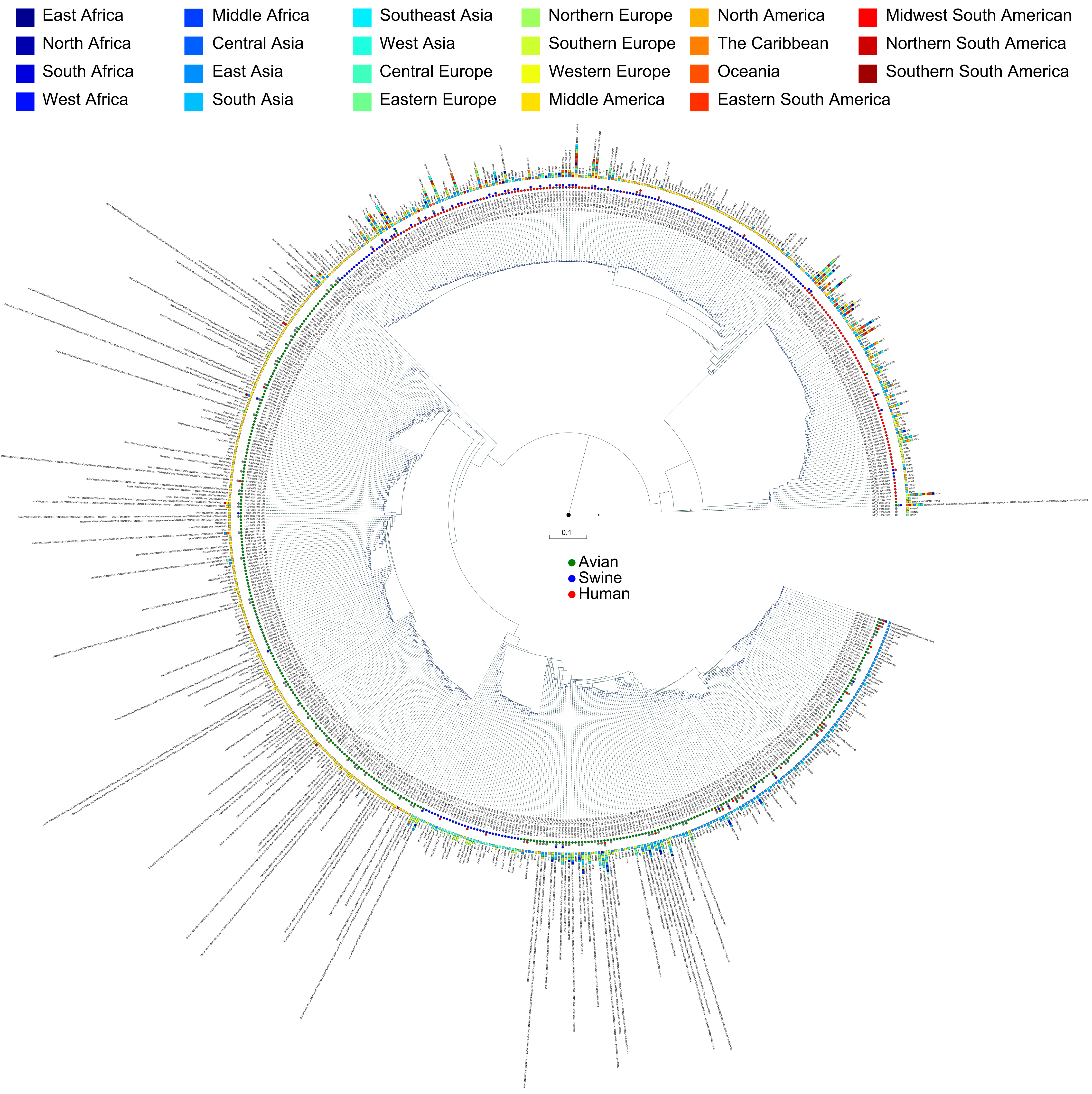
