## Supplementary Figures for "Reassortment Network of Influenza A Virus": Supplementary Figure SF5.pdf

Supplementary Figure SF5. The detailed type result for NA segment. The year range, hosts, locations and subtypes are shown after each NA type, where the circles represent hosts and the rectangles represent locations. The hosts and locations are distinguished by different colors.

East Africa, North Africa, South Africa, West Africa, Middle Africa, Central Asia, East Asia, South Asia, Southeast Asia, West Asia, Central Europe, Eastern Europe, Northern Europe, Southern Europe, Western Europe, Middle America, North America, The Caribbean, Oceania, Eastern South America, Midwest South American, Northern South America, Southern South America

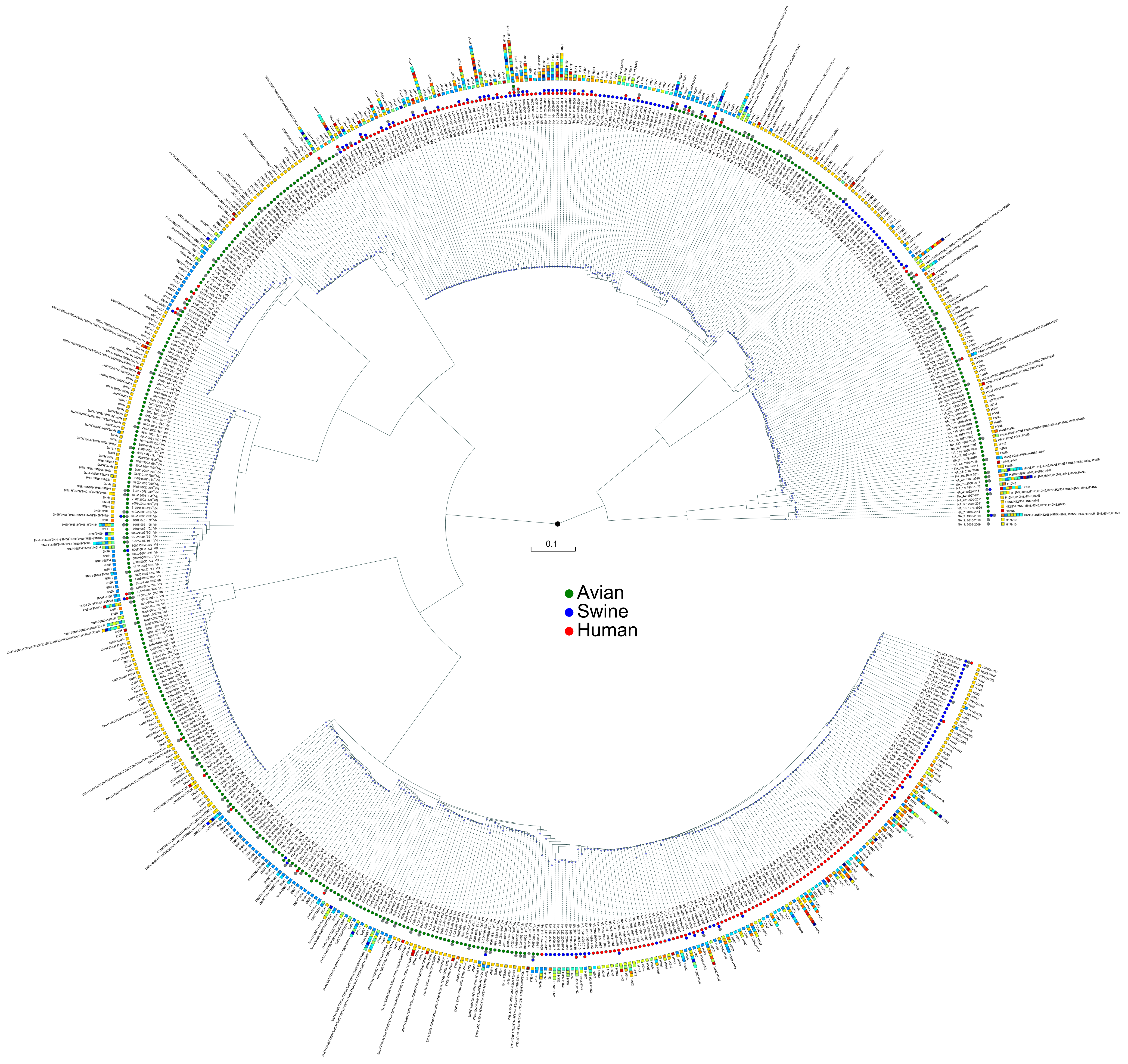
