## Supplementary Figures for "Reassortment Network of Influenza A Virus": Supplementary Figure SF6.pdf

Supplementary Figure SF6. The detailed type result for MP segment. The year range, hosts, locations and subtypes are shown after each MP type, where the circles represent hosts and the rectangles represent locations. The hosts and locations are distinguished by different colors.

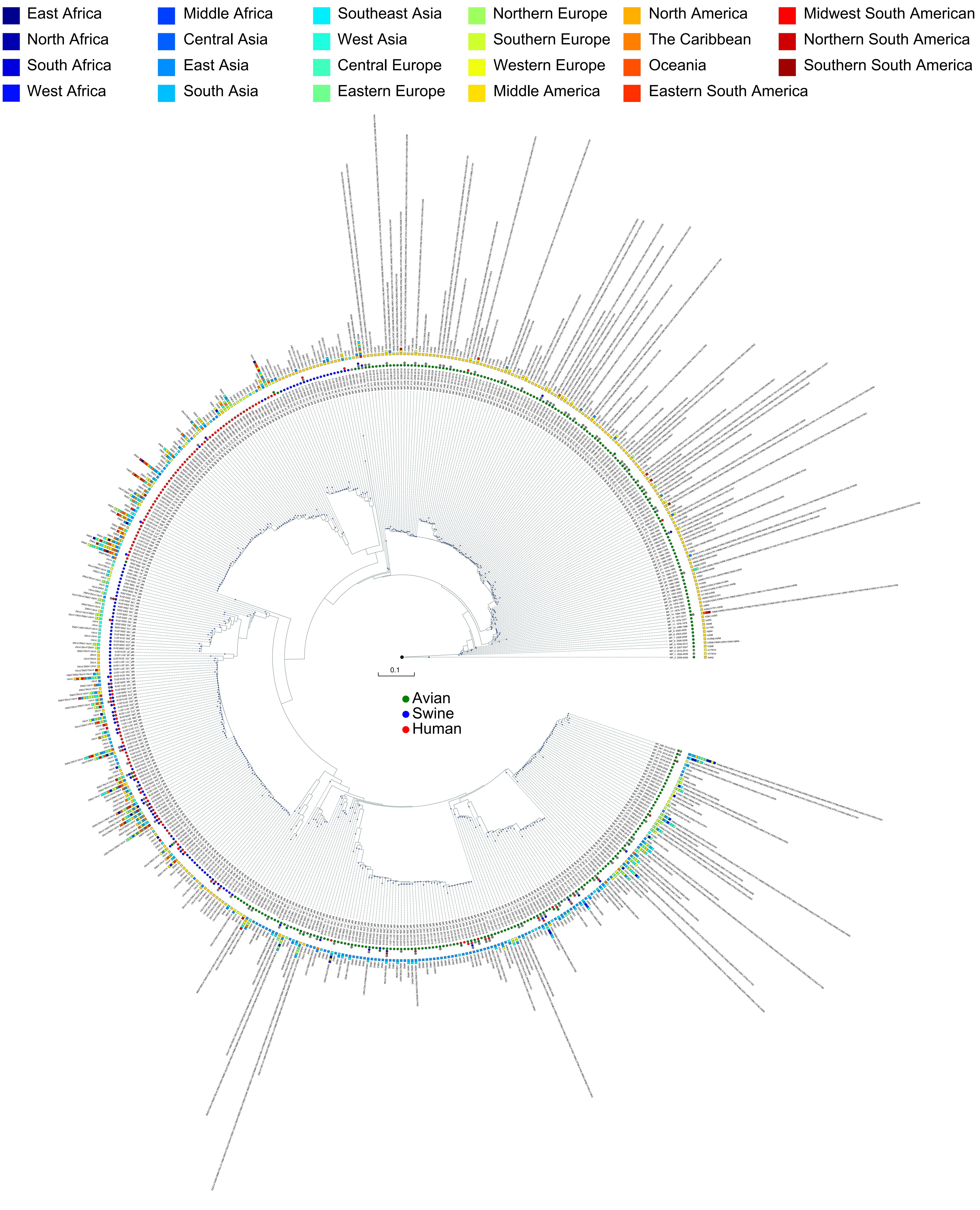
