## Supplementary Figures for "Reassortment Network of Influenza A Virus": Supplementary Figure SF8.pdf

Supplementary Figure SF8. The reassortment history of IAVs. Each sector represents a location, while the polar axis represents the years. Each circle represents a virus, with different colors to indicate the hosts. The line from source to target represents the parental virus produces the reassortant virus. Intra-Locations and Inter-Locations reassortment are indicated by green and red lines, respectively.

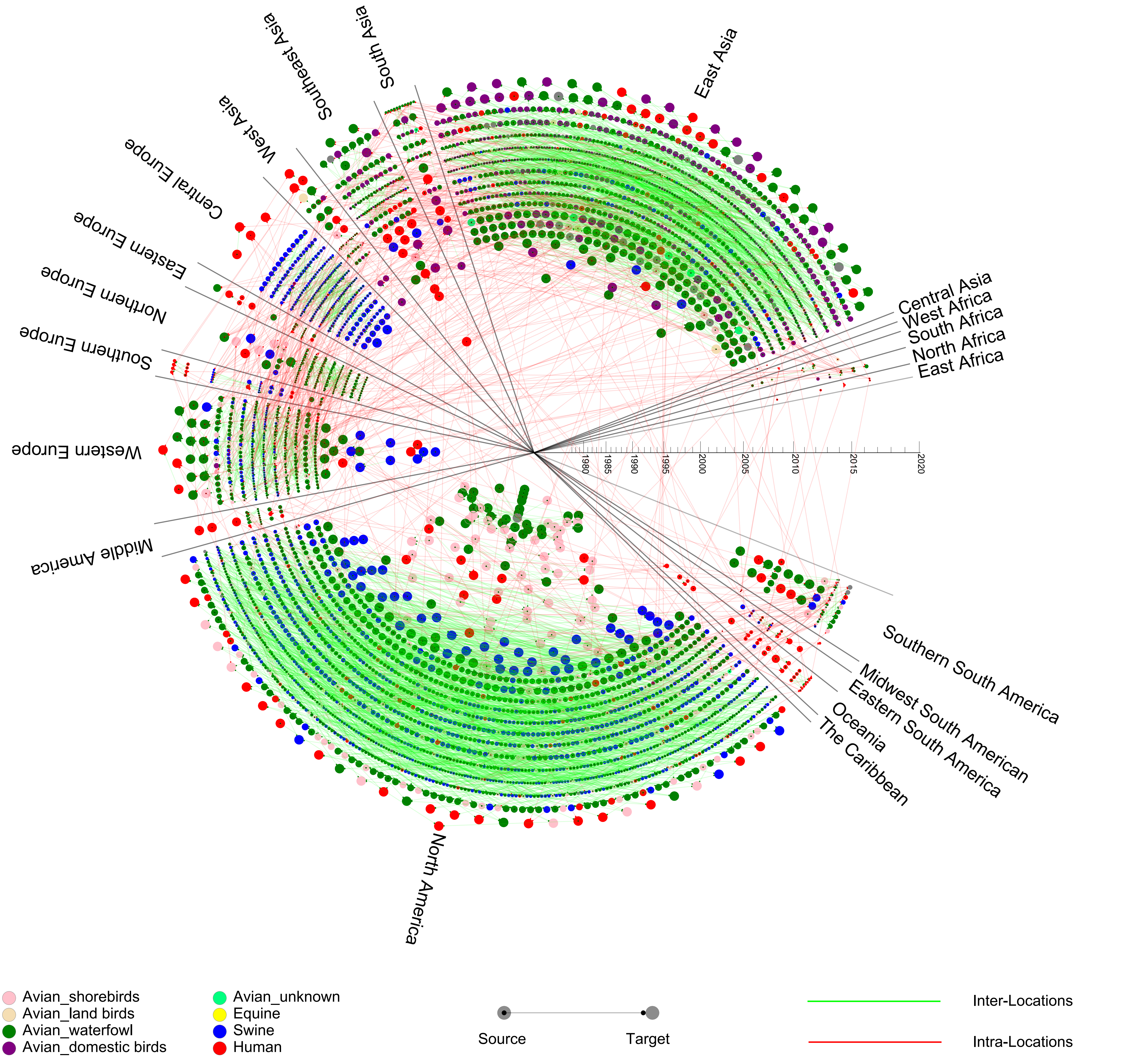
