## Supplementary Figures for "Reassortment Network of Influenza A Virus": Supplementary Figure SF9.pdf

Supplementary Figure SF9. The gene flow network of IAVs (with subtype information annotated). The subtypes of the types are distinguished by different colors, where red represents H9N2, yellow represent H7N9, green represent H1N1, blue represent H3N2, etc.

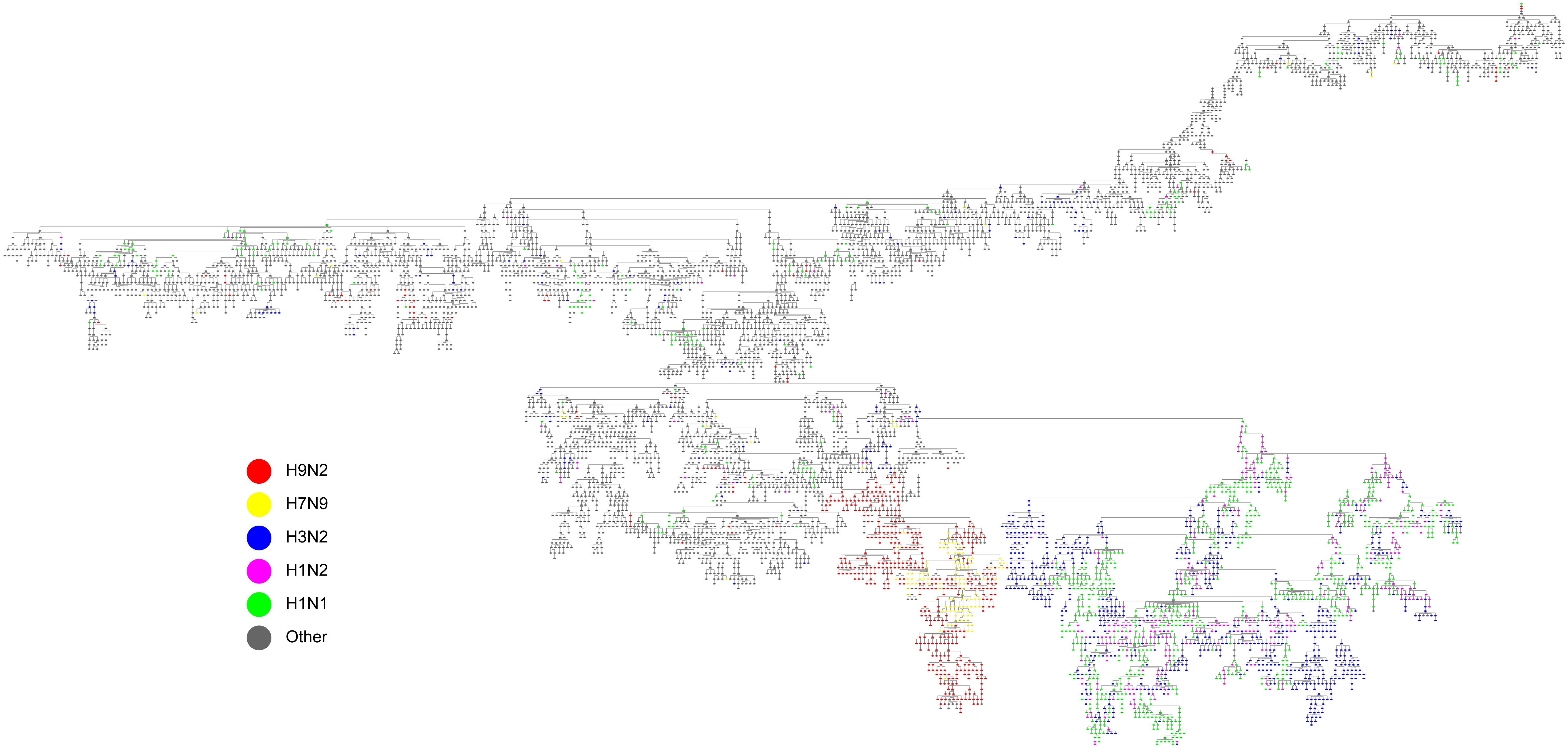
