## Supplementary Figures for "Reassortment Network of Influenza A Virus": Supplementary Figure SF10.pdf

Supplementary Figure SF10. The gene flow network of IAVs (with location information annotated). Each node is a genotype of IAVs and the edge represents gene segment flow by reassortment between two genotypes. The locations are distinguished by different colors.

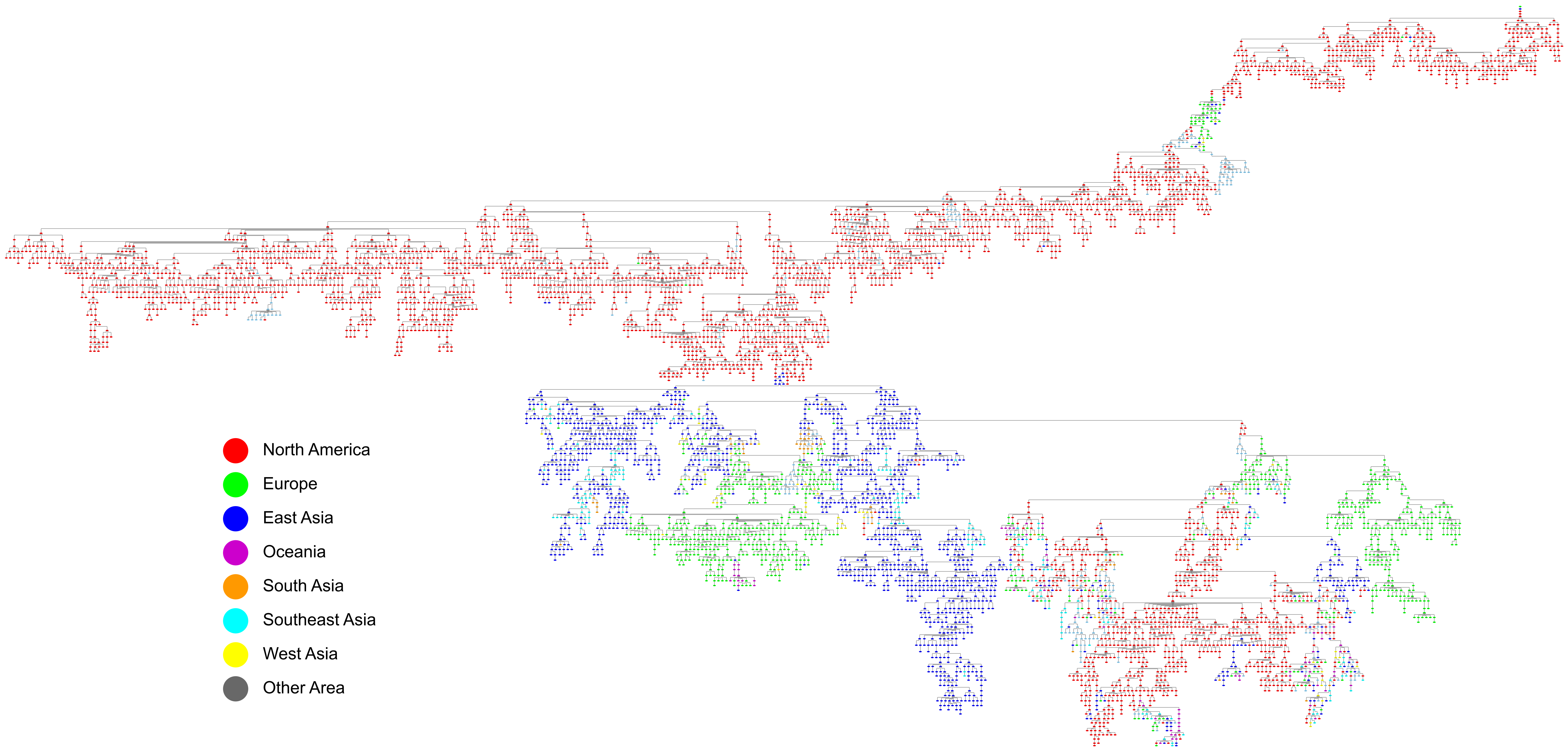
