## Supplementary Figures for "Reassortment Network of Influenza A Virus": Supplementary Figure SF11.pdf

Supplementary Figure SF11. Comparison of clustering results with unified nomenclature system. The inner circle was the clustering result of the unified nomenclature system, and the outer circle was our clustering result. Each HA segment of the virus corresponded to a small sector. We used different colors to distinguish different clades. Because of the large number of clades, the colors were used repeatedly.

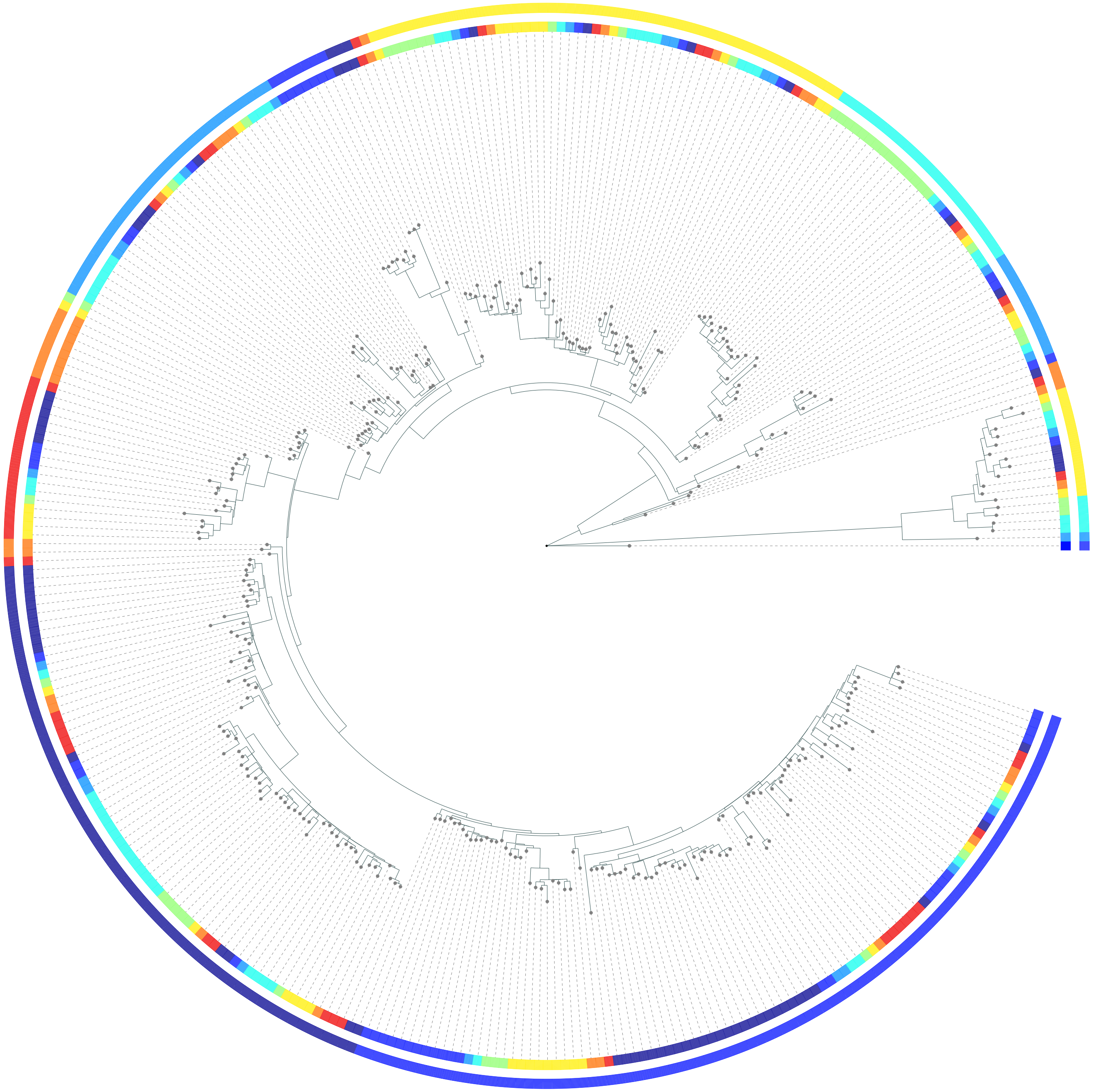
